# Manipulating Histone Acetylation Leads to Adverse Effects in Hemangiosarcoma Cells

**DOI:** 10.1101/2021.12.10.472173

**Authors:** Tamami Suzuki, Keisuke Aoshima, Jumpei Yamazaki, Atsushi Kobayashi, Takashi Kimura

**Affiliations:** Laboratory of Comparative Pathology, Department of Clinical Sciences, Faculty of Veterinary Medicine, Hokkaido University, Sapporo, Hokkaido, 060-0818, Japan; Translational Research Unit, Veterinary Teaching Hospital, Faculty of Veterinary Medicine, Hokkaido University, Sapporo, Hokkaido, 060-0818, Japan

**Keywords:** BETi, HDACi, hemangiosarcoma, histone acetylation

## Abstract

Canine hemangiosarcoma (HSA) is a malignant tumour derived from endothelial cells. No effective treatment has yet been developed because of the lack of understanding of its pathogenesis. Histone acetylation, an epigenetic modification, is highly associated with cancer pathogenesis. Manipulating histone acetylation by histone deacetylase inhibitors (HDACi) or bromodomain and extraterminal domain inhibitors (BETi) is one approach to treat various cancers. However, the role of histone acetylation in HSA remains unknown. This study aimed to investigate how histone acetylation functions in HSA pathogenesis using two HDACi, suberanilohydroxamic acid (SAHA) and valproic acid (VPA), and one BETi, JQ1, *in vitro* and *in vivo*. Histone acetylation levels were high in cell lines and heterogeneous in clinical cases. SAHA and JQ1 induced apoptosis in HSA cell lines. SAHA and VPA treatment in HSA cell lines upregulated inflammatory-related genes, thereby attracting macrophages. This implies that SAHA and VPA can induce anti-tumour immunity. JQ1 stimulated autophagy and inhibited the cell cycle. Finally, JQ1 suppressed HSA tumour cell proliferation *in vivo*. These results suggest that HDACi and BETi can be alternative drugs for HSA treatment. Although further research is required, this study provides useful insights for developing new treatments for HSA.

## Introduction

Canine hemangiosarcoma (HSA) is a malignant tumour derived from endothelial cells. Surgery and chemotherapy are conventional therapies for HSA, but the efficacy of these treatments was limited (Clifford et al., 2000). Since little is known about the molecular pathogenesis of HSA, more basic research on HSA is warranted.

Epigenetics such as DNA methylation and histone modifications regulates transcription, and it is involved in various biological processes such as embryonic development, metabolism, and diseases, including cancer (Kaelin and McKnight, 2013; Virani et al., 2012; Xu et al., 2021). Acetylated histone can promote transcription by binding to bromodomain and extraterminal (BET) proteins and loosening the chromatin structure by neutralizing the electric charges (Josling et al., 2012; Clayton et al., 2006).

In recent years, a growing number of reports have highlighted that histone acetylation is involved in cancer pathogenesis (Calcagno et al., 2019; Liu et al., 2018). Histone deacetylase inhibitors (HDACi) and BET inhibitors (BETi) have been used to investigate the function of histone acetylation and their treatment efficacy. HDACi inhibits histone deacetylase function, thereby increasing histone acetylation levels (West and Johnstone, 2014). BETi binds to the functional domain of BET proteins through which BET proteins cannot recognise acetylated histones (Bechter and Schöffski, 2020). Suberanilohydroxamic acid (SAHA) and valproic acid (VPA) are HDACis that have been reported to induce apoptosis *in vitro* and suppress tumour proliferation *in vivo* in human cancers (Claerhout et al., 2011; Lee et al., 2012; Uehara et al., 2012; Yagi et al., 2010). HDACi can also improve tumour immunity, and combination therapies with immune checkpoint inhibitors or chemotherapeutics enhance their efficacy in human cancers (Huang et al., 2017; Kim et al., 2020; Xue et al., 2016). BETi has also been broadly investigated as cancer drugs, including JQ1, which causes apoptosis, cell cycle arrest, and autophagy *in vitro*, and impede tumour growth *in vivo* (Jostes et al., 2017; Li et al., 2019; Wen et al., 2019).

Regarding epigenetics in HSA, KDM2B—a histone demethylase—has recently been found to promote tumour viability (Gulay et al., 2021). However, the role of histone acetylation in HSA remains unknown. In this study, we aimed to investigate the role of histone acetylation and the effects of HDACi and BETi on HSA.

## Materials and Methods

### Histopathology and immunohistochemistry (IHC)

Tumour samples were obtained from patients presenting to Hokkaido University Veterinary Teaching Hospital with written informed consent (Table S1). Haematoxylin and eosin staining was performed as previously described (Maharani et al., 2018). IHC analysis was performed as previously described (Gulay et al., 2021) with slight modifications. Antigen retrieval was performed in Tris-EDTA buffer (pH 9.0) in a microwave for 10 min. Tissues were stained with anti-acetylated histone H3 antibody (1:250; #39140, Active Motif, CA, USA) overnight at 4°C. Quantification of IHC results was performed with QuPath using Nucleus DAB OD mean values, and the values in tumour cells were normalised against those in normal endothelial cells on the same slides for further analysis (Bankhead et al., 2017).

### Cell culture

Canine aortic endothelial cells (CnAOEC: #Cn304-05, Cell Applications, CA, USA), seven canine HSA cell lines (JuA1, JuB2, JuB4, Re12, Re21, Ud2, and Ud6). (Murai et al., 2012), ISOS-1 obtained from the Cell Resource Center for Biomedical Research Cell Bank (Tohoku University) (Masuzawa et al., 1998), and RAW264 obtained from RIKEN Bioresource Center were cultured with Dulbecco’s Modified Eagle Medium High Glucose (DMEM; #044-29765, Fujifilm Wako Pure Chemical Industries, Osaka, Japan) supplemented with 10% foetal bovine serum (FBS; #S1580-500, Biowest, UT, USA) and penicillin– streptomycin solution (#168-23191, Fujifilm Wako) at 37□ with 5% CO_2_.

### Drug preparation

SAHA (#10009929, Cayman Chemical, MI, USA) was dissolved in dimethyl sulfoxide (DMSO) to a stock solution of 150 mg/ml. For *in vivo* studies, the stock solution of SAHA was diluted with 30% polyethylene glycol 400 and 5% Tween 80. VPA (#227-01071, Fujifilm Wako) was dissolved in PBS. JQ1 (#T117580, Toronto Research Chemicals, Toronto, Canada) was initially dissolved in DMSO to a stock solution of 50 mg/ml. For *in vivo* studies, the stock solution of JQ1 was diluted with 5% Tween 80.

### Cell viability assay

Two thousand cells were seeded in 96-well cell culture plates. The next day, they were treated with DMSO, SAHA, or JQ1. Survival rates and growth inhibition rates were analysed as previously described (Aoshima et al., 2018; Morita et al., 2019)^1^.

### Apoptosis assay

Apoptosis assay was performed using the FITC Annexin V Apoptosis Detection Kit [(#556547, BD Biosciences, NJ, USA) according to the manufacturer’s instructions. Briefly, JuB4 and Re12 cells were treated with 10 μM SAHA or JQ1 for 24 h, and detached from dishes using TrypLE Express Enzyme (#12604013; Thermo Fisher Scientific). One million cells were suspended in the binding buffer and stained with annexin V. Cells were analysed with a BD FACSVerse flow cytometer (BD Biosciences), and the results were analysed with FCS Express 4 (De Novo Software, CA, USA).

### RNA-Sequencing

ISOS-1 were treated with 1 μM SAHA for 24 h, 1 μM JQ1 for 36 h, or 2 mM VPA for 36 h in triplicate. Total RNA was extracted with a NucleoSpin RNA isolation kit (#740955.50, Macherey-Nagel GmbH & Co. Düren, Germany) according to the manufacturer’s instructions. RNA samples were submitted to Kazusa DNA Research Institution (Chiba, Japan) for further analyses. RNA-seq libraries were constructed using a QuantSeq 3’mRNA-Seq Library Prep Kit (LEXOGEN, Vienna, Austria) and sequenced with the NextSeq500 (Illumina, CA, USA) to generate a minimum of two million single-end 75-bp reads. Sequencing reads were mapped to the mm39 mouse reference genome using STAR, and expression levels were estimated using RSEM (Dobin and Gingeras, 2015; Li and Dewey, 2011). Differential expression and gene expression profiles were analysed by edgeR and GSEA v4.1.0 (Robinson et al., 2009; Mootha et al., 2003; Subramanian et al., 2005), respectively.

### SA-β-gal staining

SA-β-gal staining was performed using a Senescence β-Galactosidase Staining Kit (#9860, Cell Signaling Technology, MA, USA) according to the manufacturer’s instructions. Briefly, ISOS-1 was treated with DMSO or 10 μM JQ1 for 24 h and fixed with 4% paraformaldehyde (PFA) for 10 min at room temperature (RT). Cells were stained with β-Galactosidase Staining Solution containing 5-bromo-4-chloro-3-indolyl β-D-galactopyranoside (X-Gal: #16495, Cayman Chemical) for 13 h at 37□. Positive cells were counted manually using an inverted microscope (BIOREVO BZ-9000; KEYENCE, Tokyo, Japan).

### Cell cycle analysis

ISOS-1, JuB2, JuB4, and Re12 were treated with DMSO or JQ1 for 24 h. ISOS-1 was treated with 10 μM JQ1, and other cell lines were treated with 5 μM JQ1. Cell cycle analysis was performed as previously described (Gulay et al., 2021).

### Migration assay

JuB4 and Re12 were treated with DMSO, 2 μM SAHA, or 2 mM VPA for 48 h in 6-well plates. Then, after replacing the medium to the one without inhibitors, the cells were co-cultured with RAW264 seeded on ThinCert Cell Culture Inserts (#657630, Greiner Bio-One) for 24 h. RAW264 on the culture surface of the inserts were removed with a cotton swab. Migrated RAW264 on the bottom side of the inserts were fixed with 4% PFA for 30 min at RT and then stained with 0.01% crystal violet for 30 min at RT. The number of cells was counted manually in 10 fields at 200× under a light microscope (BH-2; Olympus, Tokyo, Japan).

### Western blotting

Protein extraction and western blotting were performed as previously described (Gulay et al., 2021). Antibodies used in this study are listed in Table S2.

### Reverse transcription quantitative polymerase chain reaction (RT-qPCR)

Primers and samples were prepared as described previously (Gulay et al., 2021). RT-qPCR was performed using the primers listed in Table S3. Results were normalised according to the geometric mean of reference genes (*RPL32* and *HPRT* for canine genes and *Hprt* and *Tbp* for murine genes), which were selected from potential internal controls by geNorm (Fig. S1) (Vandesompele et al., 2002). Relative expression levels were calculated by setting the DMSO-treated cells as the control.

### Animal studies

All mouse experiments were performed under the guidelines of Hokkaido University (protocol number: 20-0083), which follows the ARRIVE guidelines. Five-week-old female Balb/c and KSN/Slc mice (Japan SLC, Inc. Shizuoka, Japan) were used. A day before tumour cell inoculation, KSN/Slc mice were treated with 100 μL 2.5 mg/ml anti-asialo GM1 (014-09801, Fujifilm Wako) to increase the success rate of JuB2 transplantation (Yoshino et al., 2000). Five million cells were inoculated subcutaneously in both flanks of Balb/c and KSN/Slc mice, which were anesthetised with 0.3 mg/kg medetomidine (Domitor, ZENOAQ, Tokyo, Japan), 4 mg/kg midazolam (Dormicum, Maruishi Pharmaceutical Co., Ltd. Osaka, Japan) and 5 mg/kg butorphanol (Vetorphale, Meiji Seika Pharma Co., Ltd. Tokyo, Japan). After tumour cell inoculation, mice were awoken by intraperitoneal injection of 3 mg/kg atipamezole (Atipame, Kyoritsu Seiyaku Corporation, Tokyo, Japan). Tumour volumes were calculated using the formula: Volume = (Length × Width^2^)/2. When the tumour volume reached 100 mm^3^, mice were treated with the vehicle control or inhibitors. Mice were intraperitoneally injected with SAHA (150 mg/kg) daily, VPA (200 mg/kg) five times weekly, or JQ1 (50 mg/kg) daily. Mice were examined at least twice weekly to check their health status and to measure tumour sizes. Balb/c mice were euthanized with CO_2_ when tumours reached 1000 mm^3^ in volume. KSN/Slc mice were euthanized with CO_2_ 11 or 12 days after treatment initiation.

### Statistical analysis

Statistical analyses were performed with R (version 4.1.0). Fisher’s exact test was used to analyse the ratio of acetylated H3 staining levels. Student’s *t*-test or Mann-whiteny test were used to analyse the differences between two groups, whereas Tukey’s test was used to analyse differences among multiple groups. Survival curves were analysed using the log-rank test. Overall tumour growth was analysed with two-way ANOVA.

## Results

### Histone acetylation levels in HSA cells are high in vitro and heterogeneous in vivo

We first analysed histone acetylation levels in HSA cell lines and CnAOEC. We found that global acetylation of H2B, H3, and H4 was significantly enriched in all HSA cell lines compared with CnAOEC (Fig. 1A). Then, we examined global acetylated histone H3 levels in 10 clinical HSA cases. These cases were classified into three types according to their histological pattern: solid, capillary, or cavernous (Kim et al., 2015). Five cases had a single histological pattern, while the other five possessed multiple proliferation patterns (Fig, 1B). We also classified histone H3 acetylation levels into three groups based on the normalized values: negative (< 0.5), weak (0.5–2.0), and strong (> 2.0) (Figs. 1C and 1D). The results showed that the histone H3 acetylation levels of tumour cells were heterogeneous, and the ratio was different depending on their histological pattern even in the same tumour tissue (Figs. 1E and 1F). Furthermore, the ratio of tumour cells classified as negative was the highest in the solid pattern followed by the capillary pattern, and the lowest in the cavernous pattern (Fig. 1F). These results suggest that histone H3 acetylation levels are heterogeneous in tumour cells and are associated with their histopathological patterns.

**Figure 1.**
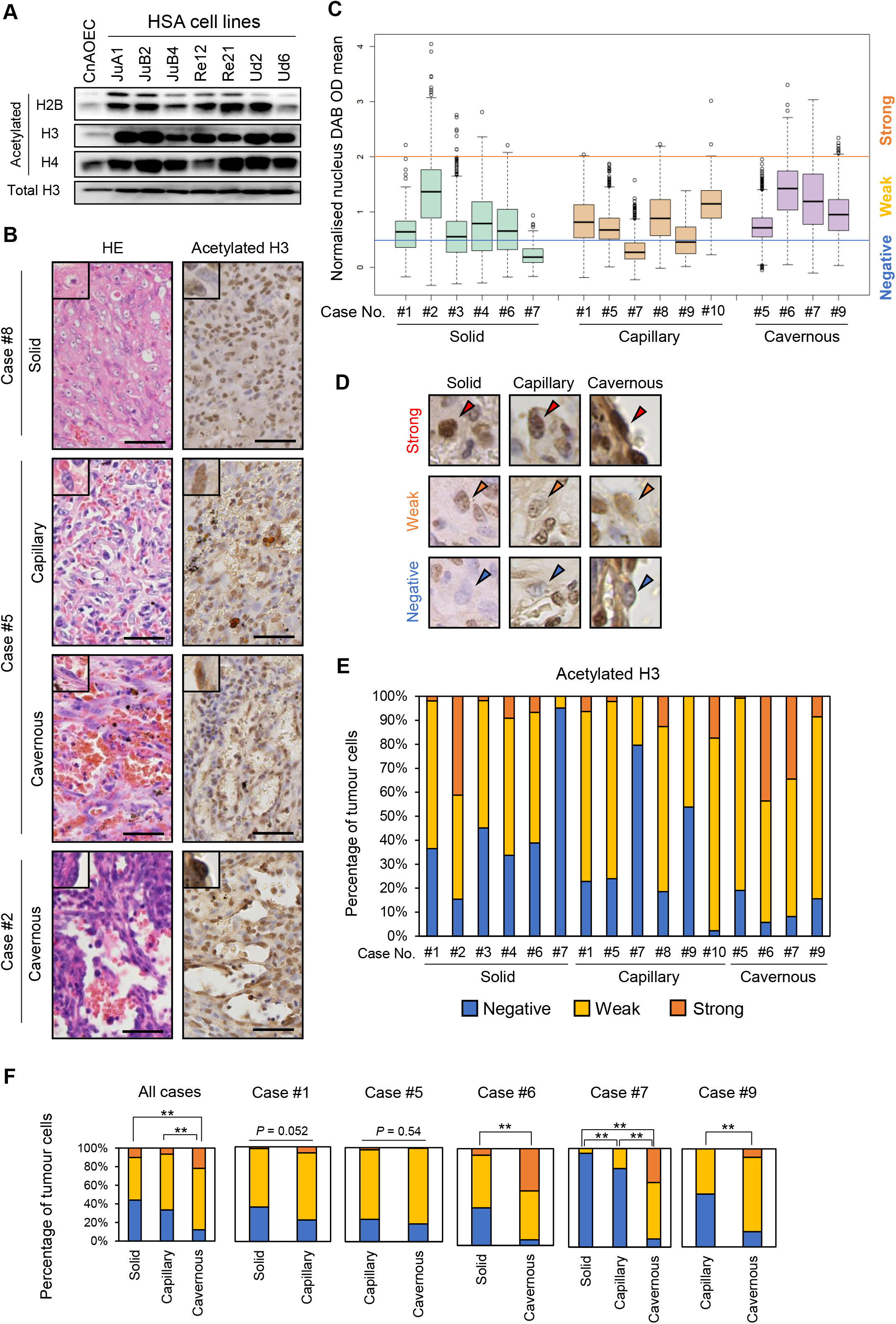
Histone acetylation levels in HSA cells are high *in vitro* and heterogenous *in vivo*. (A) Western blot analysis of acetylated histone H2B, H3, and H4. (B) Representative images of HE and IHC staining in clinical HSA cases. Scale bars = 50 μm. (C) Box plots for normalized nucleus DAB OD mean values in 10 clinical HSA cases. (D) Representative IHC images of the classifications according to normalised acetylated H3 intensities in each histological pattern. Arrowheads indicate the nuclei of tumour cells. (E) Stacked bar graphs indicating the ratio of each staining level in 10 clinical cases. (F) Stacked bar graphs comparing the ratio of each staining level according to the histological patterns in all cases or within each case. \**P* < 0.05, \*\**P* < 0.01. Fisher’s exact test.

### SAHA and JQ1 induce apoptosis in HSA cell lines

To examine the roles of histone acetylation in HSA, we modified histone acetylation levels using HDACi (SAHA and VPA) and BETi (JQ1). First, we evaluated whether global histone acetylation levels were altered by these inhibitors. SAHA and VPA treatments increased global histone acetylation levels in both canine and murine HSA cell lines, whereas JQ1 decreased them in canine HSA cell lines but not in ISOS-1 (Fig. 2A). Then, we examined the effects of HDACi and BETi on HSA cell proliferation. SAHA and JQ1 treatment significantly decreased the number of cells, while VPA treatment did not affect their proliferation (Fig. 2B). IC_50_ values of the inhibitors in HSA cell lines were almost or less than one-tenth of those in CnAOEC, which indicated that HSA cell lines were more susceptible to these inhibitors than CnAOEC (Fig. 2C). Then, we checked apoptosis markers and found that SAHA and JQ1 increased cleaved caspase 3 expression levels and annexin V positivity in HSA cell lines except for JQ1-treated ISOS-1 (Figs. 2D–F). Finally, we evaluated protein expression and phosphorylation levels of cell survival proteins AKT and ERK. Although the results were different among the cell lines, either total or phosphorylated AKT or ERK levels were downregulated by SAHA and JQ1 but not by VPA (Fig. S2B). These results suggested that SAHA and JQ1 induced apoptosis in HSA cell lines except for ISOS-1 treated with JQ1, and that VPA did not induce cell death in all HSA cell lines.

**Figure 2.**
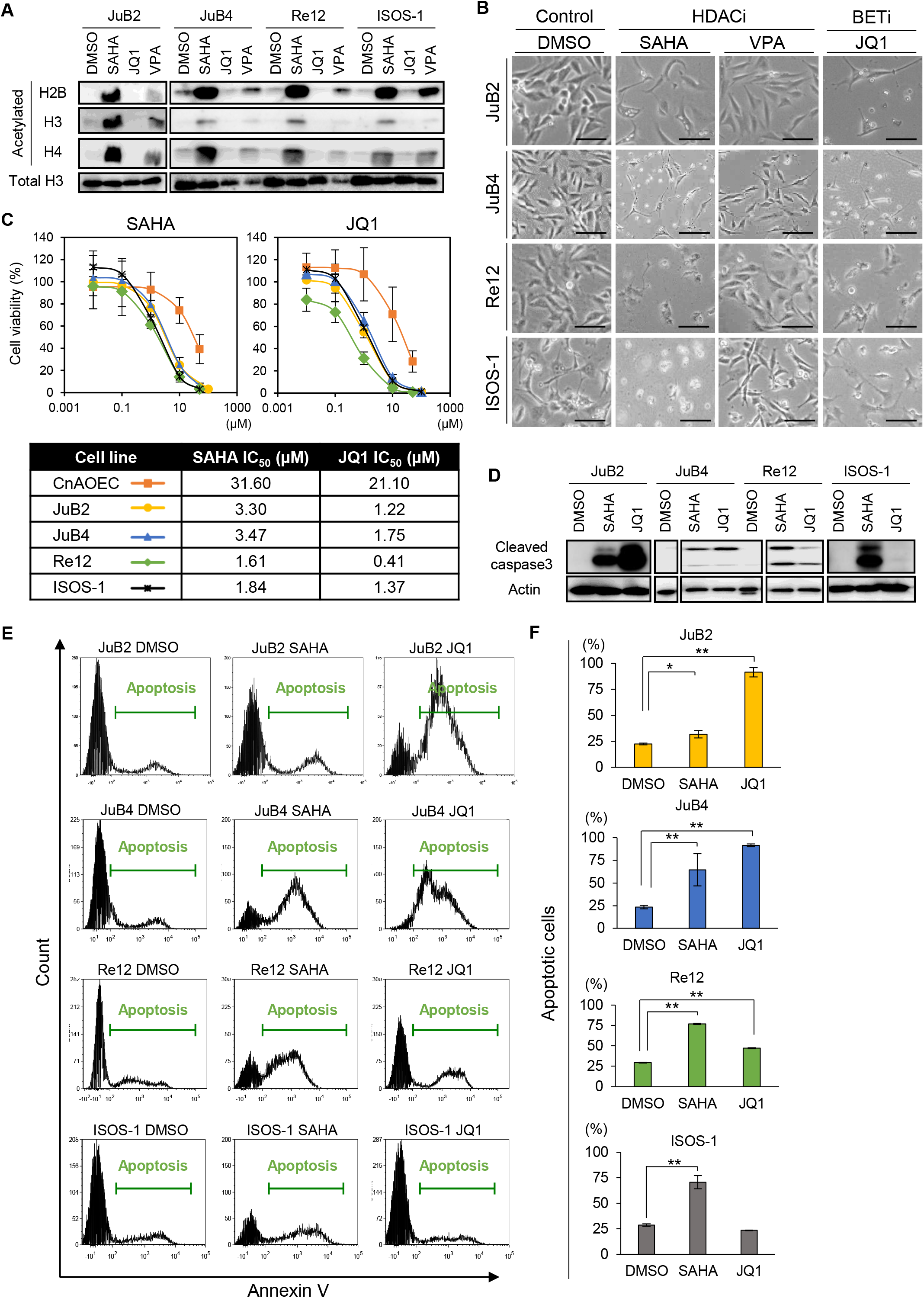
SAHA and JQ1 induce apoptosis in HSA cell lines. (A) Western blot analysis of acetylated H2B, H3, and H4 in canine HSA cell lines (JuB2, JuB4, Re12) and a murine HSA cell line (ISOS-1) treated with DMSO, SAHA, JQ1, or VPA. (B) Phase contrast images of canine and murine HSA cell lines 48 h after treatment with DMSO, SAHA, VPA, or JQ1. Scale bars = 100 μm. (C) Survival curves (top) and IC_50_ values (bottom) of canine and murine HSA cell lines treated with SAHA or JQ1 for 60 h. All samples were analysed in triplicate. Data are plotted as average percentages ± SD. (D) Western blot analysis of cleaved caspase 3 in canine and murine HSA cells treated with DMSO, SAHA, or JQ1. Uncropped images of JuB4 and Re12 are shown in Figure S2A. (E) Annexin V staining of canine and murine HSA cells treated with DMSO, SAHA, or JQ1. (F) Percentages of apoptotic cells (annexin V-positive) in canine and murine HSA cells treated with DMSO, SAHA, or JQ1. All samples were analysed in triplicate. Data are plotted as average percentages ± SD. \**P* < 0.05, \*\**P* < 0.01. Tukey’s *t* test.

### SAHA and VPA activate inflammatory responses in HSA cell lines

We further investigated the mechanisms of their effects in HSA cells. GSEA indicated that inflammatory-related gene expression was positively correlated in SAHA and VPA treatment in ISOS-1 (Fig. 3A). Consistent with this, *Il6, Cxcl1, Ccl2, Ccl7*, and *Oas1a* were upregulated more than two-fold following treatment with either SAHA or VPA or both (Fig. 3B). We wanted to extrapolate these findings to canine HSA, but *CXCL1* and *CCL2* do not exist in the canine genome and *CCL7* expression was not detected. However, *OAS1*, the orthologue of murine *Oas1a*, and other inflammatory-related genes were upregulated more than two-fold in canine HSA cell lines treated with SAHA and/or VPA (Fig. 3C).

**Figure 3.**
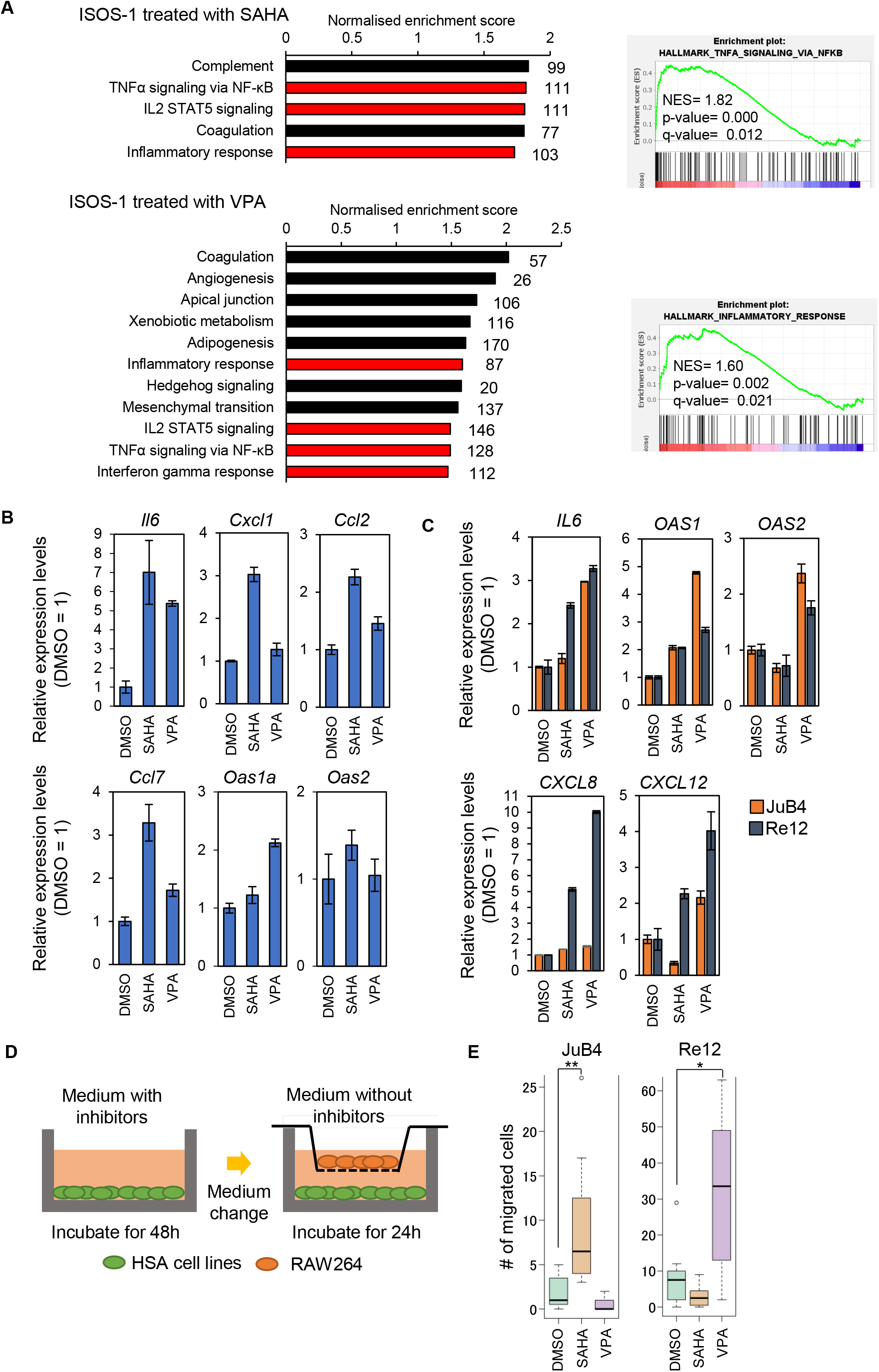
SAHA and VPA activate inflammatory responses in HSA cell lines. (A) (Left) GSEA results in ISOS-1 treated with SAHA (top) or VPA (bottom). Inflammatory-related gene sets are highlighted in red. The number of genes in each gene set is shown next to each bar. (Right) Representative GSEA plots of enriched gene sets in ISOS-1 treated with SAHA (top) or VPA (bottom). (B) RT-qPCR results of genes related to inflammatory responses in ISOS-1 treated with SAHA or VPA. (C) RT-qPCR results of inflammatory-related genes in JuB4 and Re12 treated with SAHA or VPA. (D) Graphical images of migration assay. (E) Quantitative analysis of migrated RAW264 cells co-cultured with SAHA or VPA-treated Re12. \**P* < 0.05, \*\**P* < 0.01. Tukey’s *t* test.

Inflammatory responses in tumour cells can induce immune cell migration, leading to host immune responses to tumour tissues. Therefore, to corroborate whether SAHA and VPA treatment in HSA cell lines can attract immune cells, a macrophage migration assay was performed using RAW264 (Fig. 3D). RAW264 migrated when they were co-cultured with SAHA-treated JuB4 or VPA-treated Re12 (Fig. 3E).

These results suggest that SAHA and VPA can induce inflammatory responses in HSA cell lines. This could attract macrophages, although it depends on the combination of HSA cell lines and HDACi.

### JQ1 induces autophagy and impedes the cell cycle in HSA cell lines

We also investigated the genome-wide gene expression changes in JQ1-treated ISOS-1. GSEA showed that autophagy-related gene expression and cell cycle related-gene expression were positively and negatively correlated, respectively (Figs 4A and 4B). These expression changes were validated by RT-qPCR (Fig. 4C). LC3-II, an active autophagy marker, was expressed at significantly higher levels in all JQ1-treated HSA cell lines except for Re12 compared with DMSO controls (Fig. 4D).

**Figure 4.**
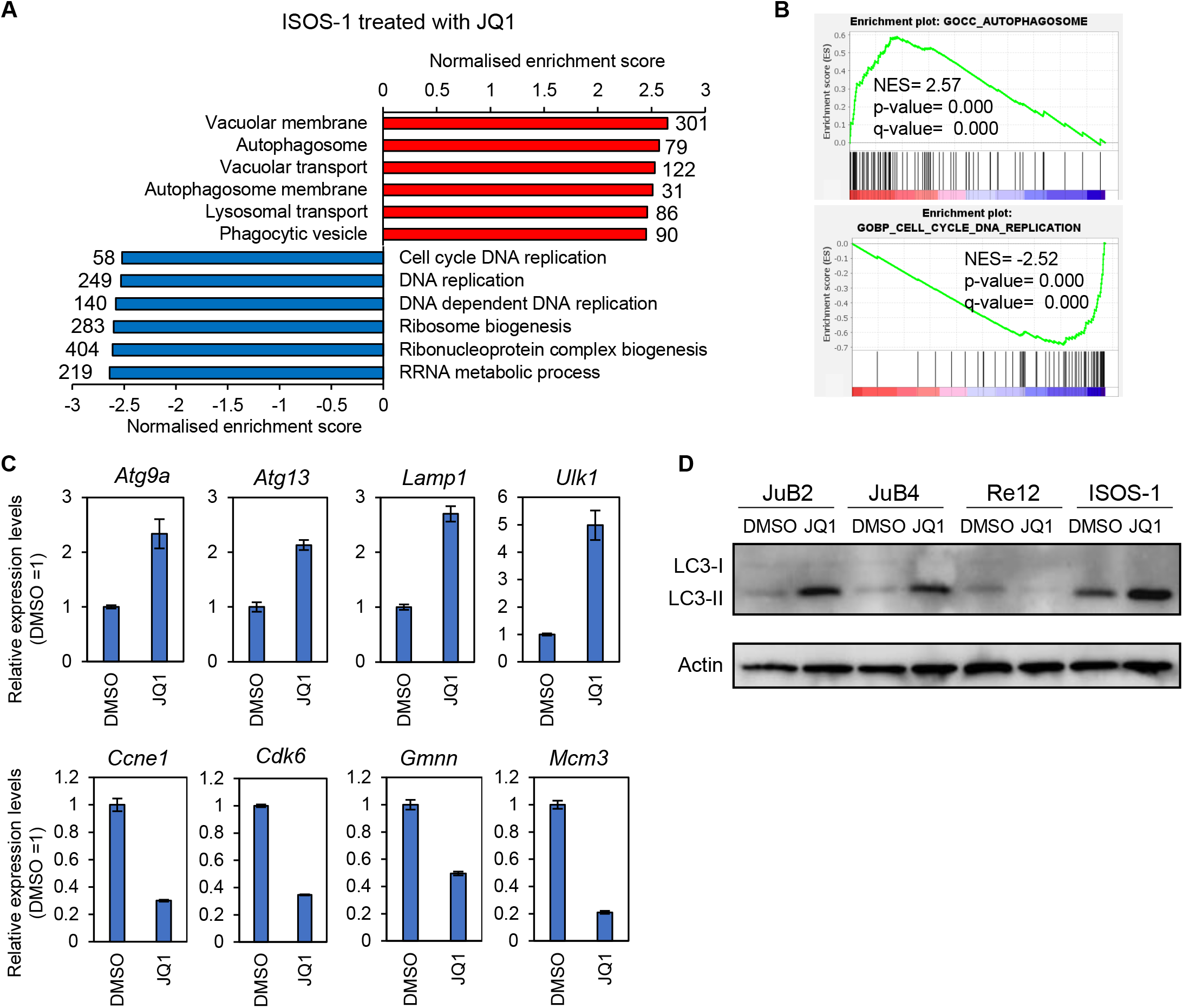
JQ1 treatment induces autophagy in canine and murine HSA cell lines. (A) Positively (top) or negatively (bottom) enriched gene sets by GSEA in ISOS-1 treated with JQ1 compared with DMSO-treated cells. The number of genes in each gene set is shown next to each bar. (B) Representative GSEA plots of positively (top) and negatively (bottom) enriched gene sets. (C) RT-qPCR of genes related to autophagy (top) and cell cycle (bottom) in ISOS-1 treated with DMSO or JQ1. (D) Western blot analysis of LC3 protein in canine and murine HSA cell lines treated with DMSO or JQ1.

Then, we generated growth inhibition curves through which we could evaluate whether the cells actively died or slowed their proliferation by JQ1. The results showed that growth inhibition rates were positive even when the cells were treated with 10 μM JQ1, which indicated that the cell cycle was impeded, and active cell death did not occur (Fig. 5A). BrdU and propidium iodide (PI) staining revealed that proliferating cells (BrdU+ cells) were significantly decreased in both murine and canine HSA cell lines by JQ1 (Fig. 5B). PI signals showed that the percentage of dead cells was slightly increased by JQ1 in ISOS-1, but its extent was significantly lower than in canine HSA cell lines (Fig. 5C). We also examined cell senescence by performing SA-β-gal staining. The results indicated that JQ1-treated ISOS-1 showed a significantly higher number of positive cells than DMSO-treated cells (Fig. 5D).

**Fig. 5.**
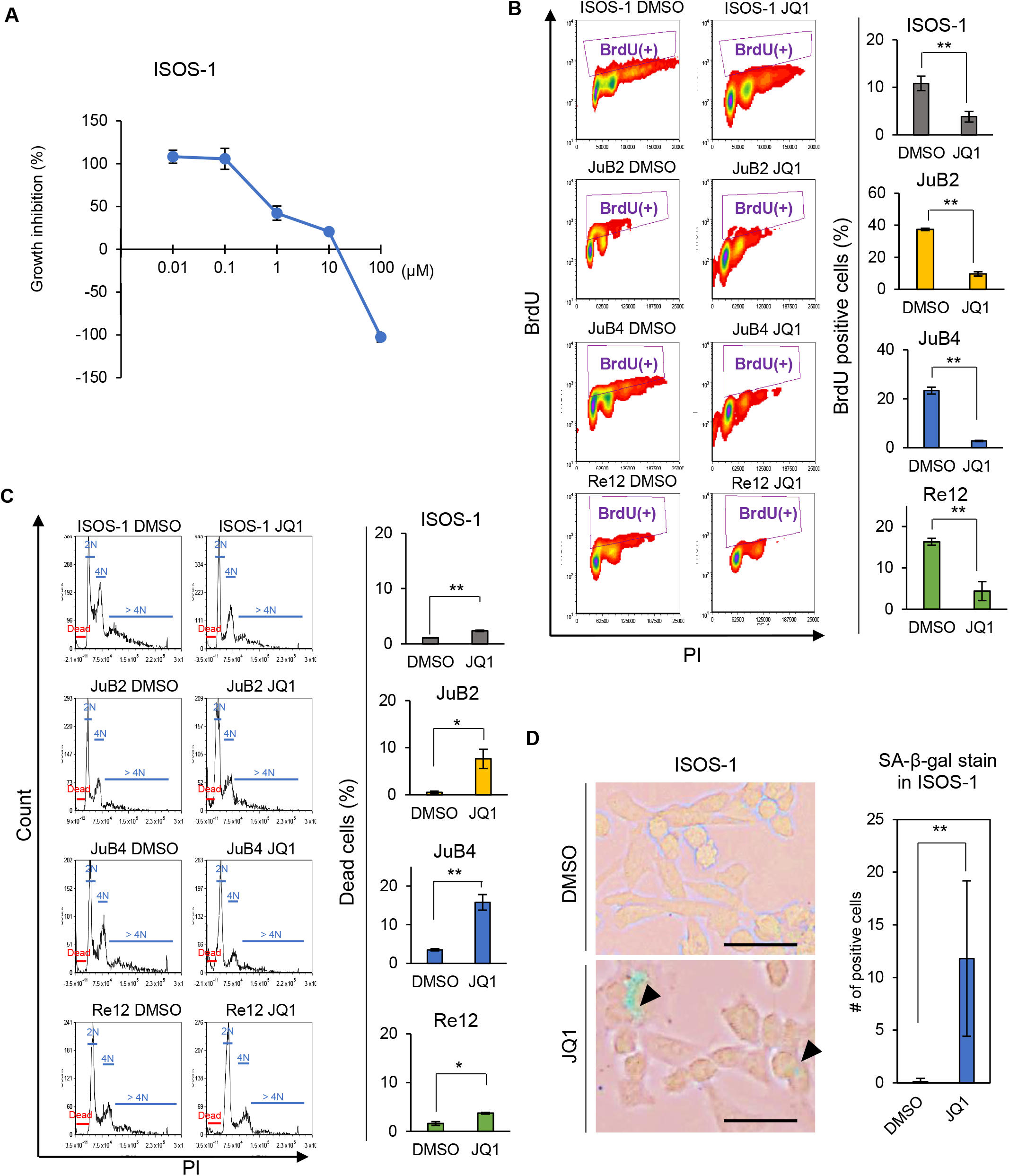
JQ1 treatment impedes the cell cycle in canine and murine HSA cell lines. (A) Growth inhibition curve of ISOS-1 treated with JQ1. All samples were analysed in triplicate. Data are plotted as average percentages ± SD. (B) Density plots (left) and quantitative analysis (right) of BrdU-positive cells in canine and murine HSA cell lines treated with DMSO or JQ1. All samples were analysed in triplicate. (C) Histograms of PI-positive cells (left) and quantitative analysis of dead cells (right) in canine and murine HAS cell lines treated with DMSO and JQ1. All samples were analysed in triplicate. (D) (Left) Representative images of SA-β-gal staining in ISOS-1 treated with DMSO or JQ1. Arrowheads indicate positive cells. (Right) Quantitative analysis of SA-β-gal-positive cells. Positive cells were counted in 10 fields under 200× magnification with a light microscope. The average numbers were plotted with ± SD. Scale bars = 100 μm. \**P* < 0.05, \*\**P* < 0.01. Student’s *t*-test.

These results suggest that autophagy activation and cell cycle downregulation are induced by JQ1, and that cell senescence but not apoptosis is caused by JQ1 in ISOS-1.

### JQ1 suppresses HSA tumour growth in vivo

Finally, to evaluate the treatment effects of SAHA, VPA, and JQ1 *in vivo*, we treated ISOS-1 tumour-bearing Balb/c mice with SAHA, VPA, or JQ1. The results showed that only JQ1 slowed tumour growth and extended the survival rates (Fig. 6A and B). We also treated JuB2 tumour-bearing KSN/Slc mice with JQ1 and showed that JQ1 also suppressed JuB2 tumour growth (Fig. 6C). These treatments did not cause body weight loss in mice (Fig. S3).

**Figure 6.**
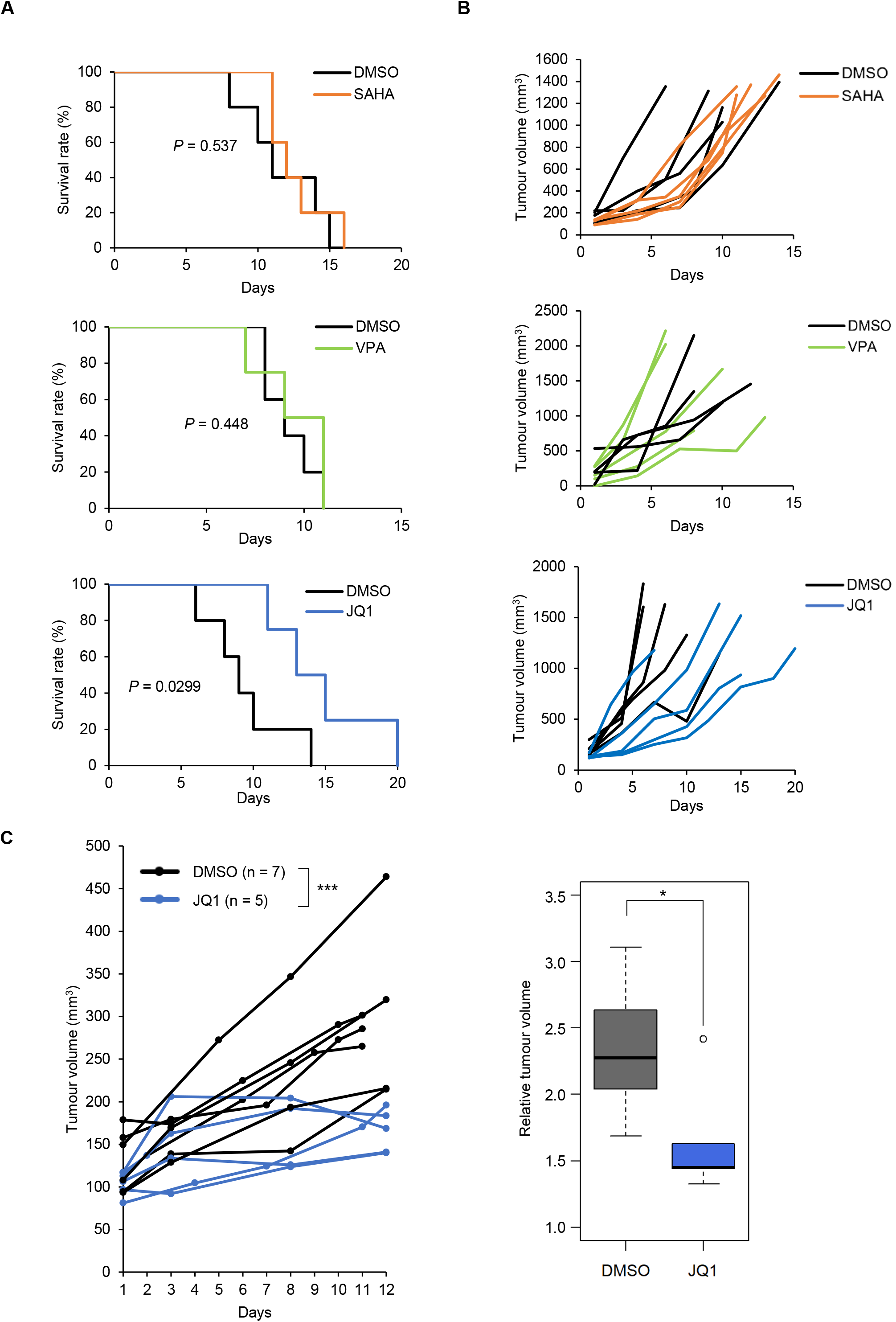
JQ1 suppresses HSA tumour growth *in vivo*. (A) Kaplan–Meier survival curves of ISOS-1-inoculated Balb/c mice treated with DMSO (n=5), SAHA (n=5), VPA (n=4), or JQ1 (n=5). Days were counted after initiating drug treatment. *P-*values were calculated using the log-rank test. (B) Individual growth curves of ISOS-1 tumours in Balb/c mice treated with DMSO, SAHA, VPA, or JQ1 after treatment initiation. (C) (Left) Individual tumour growth curves of JuB2 tumours in KSN/Slc mice treated with DMSO or JQ1. The day when the treatment began was defined as day 1. \*\*\**P* < 0.001. Two-way ANOVA. (Right) Box plots of relative tumour volumes on the last day in KSN/Slc mice treated with DMSO or JQ1. Relative tumour volumes were calculated by dividing the tumour volume on the last day by that at day 1. \**P* < 0.05, Mann-whitney U test.

These results suggest that JQ1 has suppressive effects on HSA tumour growth *in vivo*.

## Discussion

HSA tumour cells in clinical cases showed various H3 acetylation levels even in the same tumour tissue. Moreover, this heterogeneity differed depending on the histological pattern. This can be explained by the differentiation status of the tumour cells. Global histone acetylation levels have been reported to be increased as cells differentiate (Ashktorab et al., 2009; Golob et al., 2008). Furthermore, HSA tumour cells forming the solid pattern are considered to be undifferentiated (Göritz et al., 2013). Given that the percentage of tumour cells classified as H3 acetylation-negative was high in the solid pattern, they could be more undifferentiated than other tumour cells. Further research is required to clarify whether global histone acetylation levels reflect the differentiation status of HSA tumour cells.

We demonstrated that SAHA and VPA upregulated inflammatory-related genes. Moreover, SAHA-treated JuB4 and VPA-treated Re12 attracted RAW264. Several reports indicated that HDACi activated immune responses and enhanced immune cell functions (Cox et al., 2020; Laengle et al., 2020; Sun et al., 2019); thus, SAHA and VPA could also be used as alternative therapeutics for HSA. In our study, however, SAHA and VPA did not suppress tumour growth of ISOS-1 in an immunocompetent mouse model. This implies that the condition we tested was not able to sufficiently induce the immune response. Co-treatment with other drugs such as doxorubicin or immune checkpoint inhibitors would be beneficial for HSA treatment by further stimulating immune responses (Le et al., 2017; Tubbs and Nussenzweig, 2017; Igase et al., 2020; Maekawa et al., 2017). In our study, we only showed the results from the RAW264 migration assay. Further studies are required to investigate if canine-derived macrophages exhibit the same results and if the migrated macrophages possess anti-tumour functions.

We revealed that JQ1 suppressed tumour growth in HSA cell lines. Several reports demonstrated that JQ1 or other BETi had anti-tumour effects, and clinical trials are ongoing (Shorstova et al., 2021; Wen et al., 2019). Given that our results could not demonstrate tumour regression upon JQ1 treatment, further research is required to enhance its efficacy and to identify another BETi that can regress HSA tumour growth *in vivo*.

## Conclusions

We demonstrated that HDACi and BETi can be used for HSA treatment. Although further research is required, this work provides useful insights for developing new HSA treatments.

## Conflict of interest statement

The authors declare that there is no conflict of interest in this study.

## Acknowledgements

We would like to extend our sincerest gratitude to Dr. Hiroki Sakai, Gifu University, for providing canine hemangiosarcoma cell lines. We also acknowledge the efforts of Drs. Mitsuyoshi Takiguchi, Shintaro Kobayashi, and Hironobu Yasui, Faculty of Veterinary Medicine, Hokkaido University for giving useful pieces of advice and constructive discussion. We are grateful to all the members of the Laboratory of Comparative Pathology, Faculty of Veterinary Medicine, Hokkaido University for their helpful discussions, encouragement, and support. This research was supported by KAKENHI Grant-in-Aid for Young Scientists (KA, Number 20K15654) provided by Japan Society for the Promotion of Science. We thank H. Nikki March, PhD, from Edanz (https://jp.edanz.com/ac) for editing a draft of this manuscript.

## Footnotes

1 See: NCI 60 Cell Five-Dose Screen. https://dtp.cancer.gov/discovery_development/nci-60/methodology.htm (Accessed 21 October, 2021).

## Notes

### Competing Interest Statement

The authors have declared no competing interest.

