## Supplementary materials for "Manipulating Histone Acetylation Leads to Adverse Effects in Hemangiosarcoma Cells"

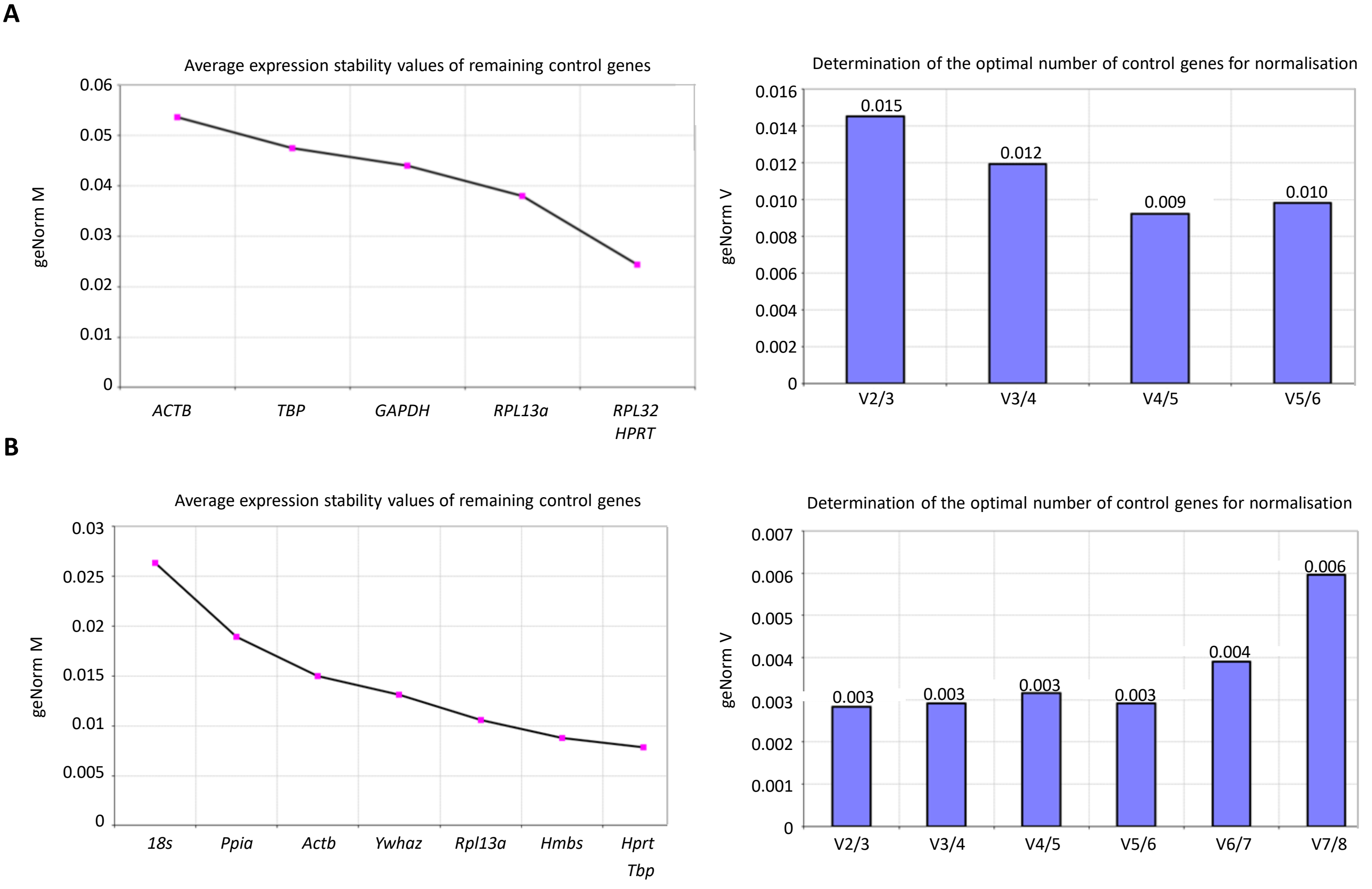


**Supplementary figure 1.**

**geNorm analysis to determine appropriate reference gene sets.**

(A, B) (Left) Average expression stability values (geNorm M) of candidate genes. The gene with the lowest M value has the most stable expression in dogs (A) and mice (B). (Right) Determination of optimal number of control genes for geometric normalisation for dogs (A) and mice (B). To determine the optimal number of reference genes, 0.15 V value was used as the cut-off value as Vandesompele et al. (2002) recommended.

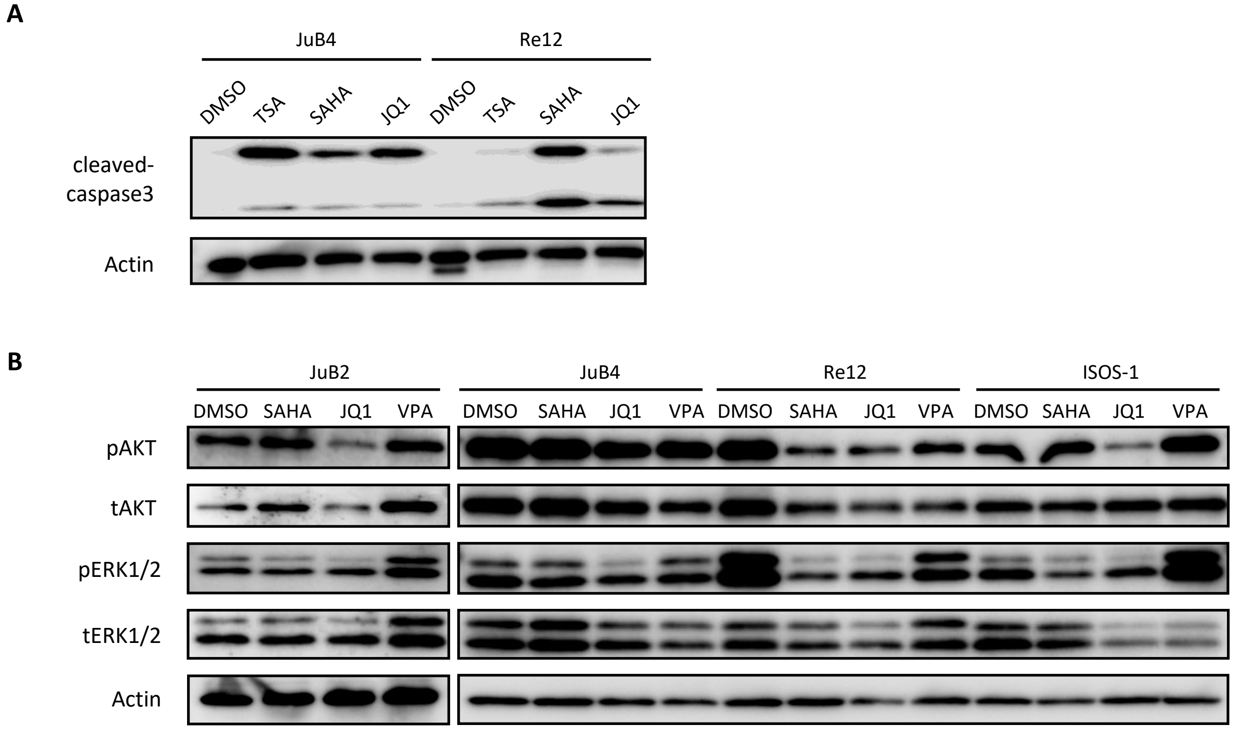


**Supplementary figure 2.**

(A) Uncropped images of western blot analysis for cleaved caspase 3 and Actin in JuB4 and Re12 treated with DMSO, TSA, SAHA and JQ1. TSA: Trichostatin A. (B) Western blot analysis for phosphorylated and total AKT and ERK expression in canine and murine HSA cell lines treated with DMSO, SAHA, JQ1 or VPA.

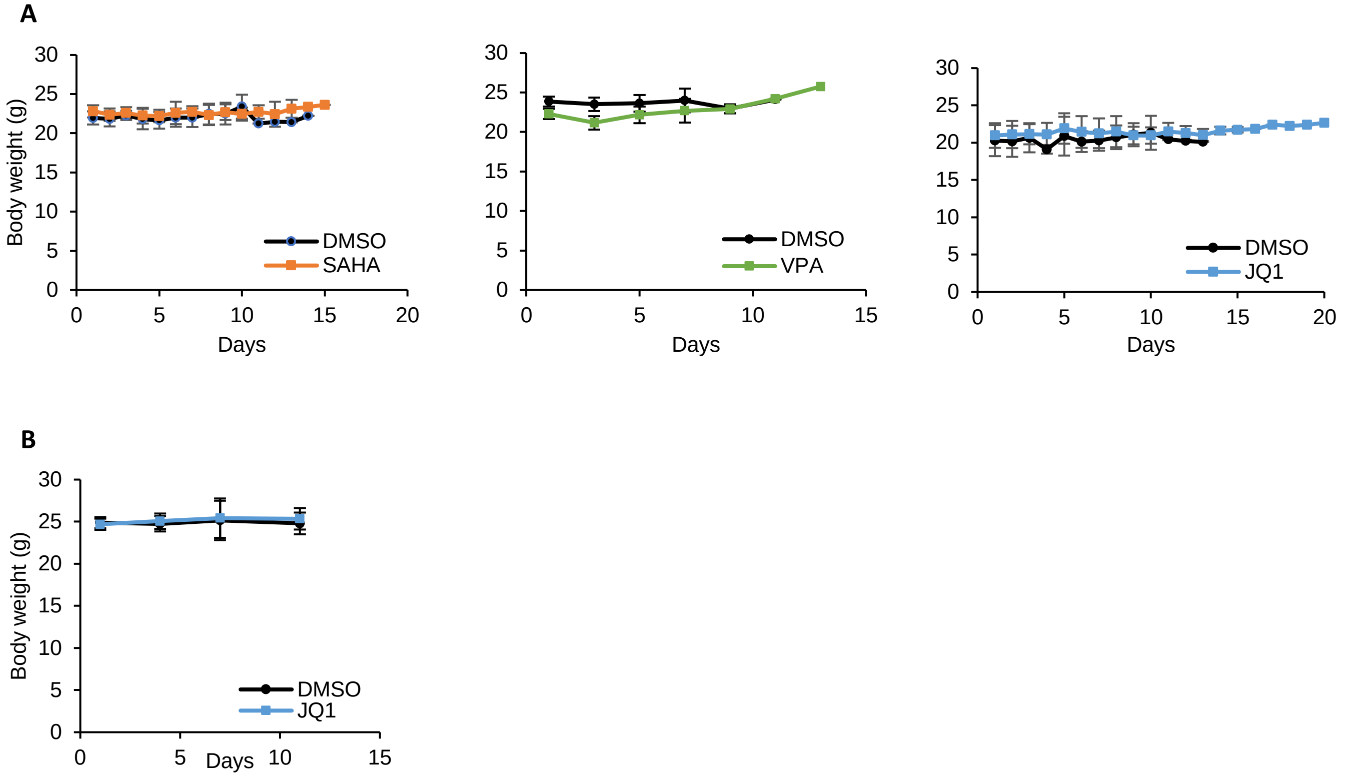


**Supplementary Figure 3.**

**Average body weight of mice in *in vivo* study**

(A) Average body weight of Balb/c mice treated with DMSO, SAHA, VPA or JQ1 after starting treatments. (B) Average body weight of KSN/Slc mice treated with DMSO or JQ1 after starting treatments.

| Case Number | Breed | Age | Sex | Location |
|---|---|---|---|---|
| 1 | Miniature Dachshund | 10y | F Spay | Spleen |
| 2 | Jack Russell Terrier | 9y | M Cast | Spleen |
| 3 | Jack Russell Terrier | 7y | F Spay | Spleen |
| 4 | Miniature Dachshund | 14y | M Cast | Spleen |
| 5 | French Bulldog | 7y | M Cast | Spleen |
| 6 | Miniature Dachshund | 9y | F | Spleen |
| 7 | Miniature Schnauzer | 11y | M | Spleen |
| 8 | Miniature Dachshund | 10y | F | Spleen |
| 9 | Flat-coated retriever | 11y | F | Spleen |
| 10 | Golden Retriever | 9y | M | Liver |

*F: female, M: male, Spay: spayed, Cast: castrated

**Supplementary Table 1.** Case information

| Protein Name | Maker | Host | Dilution | Catalog Number |
|---|---|---|---|---|
| Acetylated Histone H2B | Santa Cruz Biotechnology, Inc. | Mouse | 1:500 | sc-515937 |
| Acetylated Histone H3 | Active Motif | Rabbit | 1:5000 | 39040 |
| Acetylated Histone H4 | Santa Cruz Biotechnology, Inc. | Mouse | 1:500 | sc-377520 |
| Cleaved-caspase 3 | Cell Signaling Technology | Rabbit | 1:1000 | 9661S |
| pAKT | Cell Signaling Technology | Rabbit | 1:1000 | 4060 |
| tAKT | Cell Signaling Technology | Rabbit | 1:1000 | 4691 |
| pERK1/2 | Cell Signaling Technology | Rabbit | 1:1000 | 4370S |
| tERK1/2 | Cell Signaling Technology | Rabbit | 1:1000 | 4695S |
| LC3 | MBL | Rabbit | 1:1000 | PM036 |
| Actin | Sigma-Aldrich | Mouse | 1:10000 | MAB1501 |
| H3 | MAB Institute | Mouse | 1:10000 | MABI0001-20 |
| Goat anti-Mouse IgG (H+L) | Thermo Fisher Scientific | Goat | 1:10000 | G21040 |
| Goat anti-Rabbit IgG (H+L) | Thermo Fisher Scientific | Goat | 1:10000 | G21234 |

**Supplementary Table 2. A**ntibody list

| Species | Target | Sequence (Forward) | Sequence (Forward) | Gene ID |
|---|---|---|---|---|
| Canine | *OAS1* | TGTGCGGGTGTCTAAAGTTG | TGAACTGTCCTCGTTTCTCG | ENSCAFG 00000023556 |
| | *OAS2* | TGACCCAGATCCAGAAAACC | CCATTCGGTAGCGTCTTTTG | ENSCAFG 00000023107 |
| | *CXCL8* | GGCAGCTTTTGTCCTTTCTG | ACACTGGCATCGAAGTTCTG | ENSCAFG 00000003029 |
| | *CXCL12* | AGCCAACGTCAAGCATCTCA | TCAATGCACACCTGTCTGCT | ENSCAFG 00000007026 |
| | *IL6* | TCGGCAAAATCTCTGCACTG | TTTCTGCCAGTGCCTCTTTG | ENSCAFG 00000002733 |
| | *RPL32* | TGGTTACAGGAGCAACAAGAAA | GCACATCAGCAGCACTTCA | ENSCAFG 00000004871 |
| | *HPRT1* | CACTGGGAAAACAATGCAGA | ACAAAGTCSGGTTTATAGCCAACA | ENSCAFG 00000018870 |
| | *GAPDH* | ATTCCACGGCACAGTCAAG | TACTCAGCACCAGCATCACC | ENSCAFG 00000015077 |
| | *TBP* | ATAAGAGAGCCCCGAACCAC | TTCACATCACAGCTCCCCAC | ENSCAFG 00000004119 |
| | *RPL13A* | GCCGGAAGGTTGTAGTCGT | GGAGGAAGGCCAGGTAATTC | ENSCAFG 00000029892 |
| Mouse | *Atg9a* | ATCACCTTGGCACCACATTG | TGGTGAAGGCAACCACAAAG | ENSMUSG 00000033124 |
| | *Atg13* | ATTTGCACCCGCTCATCATC | AGGGCCTTCTTTGCTTCATG | ENSMUSG 00000027244 |
| | *Lamp1* | AGTCTTGTGTTGGCGTTCAG | AGGCAATGCATTACGTGAGC | ENSMUSG 00000031447 |
| | *Ulk1* | AAACATCGTGGCGCTGTATG | TTCACTCAGTGTGCGCATAG | ENSMUSG 00000029512 |
| | *Ccne1* | AAGCCCAAGCAAAGAAAGCC | TGGCAGGTTTGGTCATTCTG | ENSMUSG 00000002068 |
| | *Cdk6* | AAGCTGCTGACCAATTGTGC | ATACGCATGCACACACACTC | ENSMUSG 00000040274 |
| | *Gmnn* | AGGAGAACGCTGAAGATGATCC | TGCTAGCTGGTCATCCCAAAG | ENSMUSG 00000006715 |
| | *Mcm3* | CGTTCCAAGGATGTCTTTGAGC | ATGTGGCTGCCGTTTTCAAG | ENSMUSG 00000041859 |
| | *Il6* | TACCACTTCACAAGTCGGAGGC | CTGCAAGTGCATCATCGTTGTTC | ENSMUSG 00000025746 |
| | *Cxcl1* | CAAACCGAAGTCATAGCCACAC | TTTCTCCGTTACTTGGGGACAC | ENSMUSG 00000029380 |
| | *Ccl2* | TTTCCACAACCACCTCAAGC | TTAAGGCATCACAGTCCGAGTC | ENSMUSG 00000035385 |
| | *Ccl7* | AAAACCCCAACTCCAAAGCC | ACAGCGGTGAGGAATTTTGC | ENSMUSG 00000035373 |
| | *Oas1a* | AAGCACTGGTACCAACTGTG | AGGCAAAGACAGTGAGCAAC | ENSMUSG 00000052776 |
| | *Oas2* | TAGACCAGGCCGTGGATG | GTTTCCCGGCCATAGGAG | ENSMUSG 00000032690 |
| | *Hprt* | GCTTGCTGGTGAAAAGGACCTCTCGAAG | CCCTGAAGTACTCATTATAGTCAAGGGCAT | ENSMUSG 00000025630 |
| | *Tbp* | CCTTGTACCCTTCACCAATGAC | ACAGCCAAGATTCACGGTAGA | ENSMUSG 00000014767 |
| | *Ywhaz* | GAAAAGTTCTTGATCCCCAATGC | TGTGACTGGTCCACAATTCCTT | ENSMUSG 00000022285 |
| | *Actb* | AGGCCAACCGTGAAAAGATG | TGGATGGCTACGTACATGGC | ENSMUSG 00000029580 |
| | *Hmbs* | ATGAGGGTGATTCGAGTGGG | TTGTCTCCCGTGGTGGACATA | ENSMUSG 00000032126 |
| | *Rpl13a* | AGGGGCAGGTTCTGGTATTG | TGTTGATGCCTTCACAGCGT | ENSMUSG 00000074129 |
| | *Ppia* | CGCGTCTCCTTCGAGCTGTTTG | TGTAAAGTCACCACCCTGGCACAT | ENSMUSG 00000071866 |
| | *18s* | CGGCTACCACATCCAAGGAA | AGCTGGAATTACCGCGGC | ENSMUSG 00000119584 |

**Supplementary Table 3.** Primer list
